# Domain-specific DNA binding activities of BRCA1 reveal substrate preferences for homologous recombination and telomere regulation

**DOI:** 10.1101/2024.05.30.594759

**Authors:** Kaitlin Lowran, Laura Campbell, Emma Cismas, Colin G. Wu

**Author notes:** **Corresponding Author:** Colin G. Wu. K.L. and L.C. contributed equally to this paper. K.L.: Wayne State University, Detroit, MI, 48202, USA. L.C.: University of Wisconsin-Madison, Madison, WI, 53706, USA.

## Abstract

BRCA1 is a crucial mediator of homologous recombination (HR), a high-fidelity pathway for repairing double-stranded DNA breaks (DSBs) in human cells. The central region of BRCA1 protein contains two putative DNA binding domains (DBDs), yet their relative substrate specificities and functional contributions to HR remain unclear. Here, we characterized the DNA binding properties of DBD1 (amino acids 330-554), DBD2 (amino acids 894-1057), and the BRCA1 C-terminal (BRCT) repeats using biolayer interferometry. We assessed their affinities for single-stranded DNA (ssDNA), double-stranded DNA (dsDNA), and G-quadruplex (G4) structures. DBD1 exhibited the highest affinity for dsDNA, while DBD2 and BRCT bound preferentially to ssDNA and G4. These findings support a model in which DBD1 directs BRCA1 to DSB sites to facilitate DNA end resection during HR, whereas DBD2 and BRCT contribute to the role of BRCA1 in telomere maintenance and chromatin remodeling through the recognition of non-canonical DNA structures.

## INTRODUCTION

The Breast Cancer Susceptibility Gene 1 (*BRCA1*) encodes a large tumor suppressor protein that is essential for maintaining genomic integrity. BRCA1 plays pivotal roles in the repair of double-stranded DNA breaks (DSB), transcriptional regulation, and telomere maintenance (1–4). Germline mutations in *BRCA1* are associated with elevated risks of breast and ovarian cancers, as well as cardiovascular diseases (5, 6). BRCA1 facilitates DSB repair during homologous recombination (HR) by coordinating the recruitment of DNA end-resection proteins such as 53BP1 (3, 7, 8). This is subsequently followed by interaction with PALB2 to promote BRCA2/RAD51-mediated strand invasion (9–11). Additionally, BRCA1 stabilizes stalled replication forks and promotes recovery during replication stress (12, 13).

Replication stress arises from obstacles such as DNA damage, common fragile sites, and non-canonical DNA structures (14). If left unresolved, these impediments can lead to fork collapse and the formation of DSB, which must be efficiently repaired to prevent genomic instability and tumorigenesis (15, 16, 17). While multiple DSB repair pathways exist—including non-homologous end joining (NHEJ), alternative end joining, and single-strand annealing—HR is the predominant mechanism and is favored for its high fidelity (18–21).

Non-canonical DNA structures, such as D-loops, cruciform, G-quadruplexes (G4), and other fork intermediates, frequently arise during replication and transcription and must be properly resolved to maintain genomic integrity (22). BRCA1 binds to several of these structures (23–26), a function that may be relevant to its roles in telomere regulation and satellite DNA maintenance (2, 4, 27). Notably, BRCA1-deficient cells are hypersensitive to G4-stabilizing compounds such as CX-5461, which impedes replication fork progression and induces DNA breaks (28, 29).

Although BRCA1 is largely intrinsically disordered, it contains two well-characterized structured domains: an N-terminal RING domain and a C-terminal BRCT domain. The RING domain forms a heterodimer with the BARD1 tumor suppressor and functions as an E3 ubiquitin ligase (30–32). The BRCA1-BARD1 complex binds preferentially to D-loops and DNA bubbles, facilitating RAD51-mediated HR and replication fork restart at G4 regions (24, 33). In contrast, the BRCT domain acts as a phospho-protein recognition site and is prevalent in other DNA repair proteins (34). BRCA1 binds to the FANCJ helicase via the BRCT when FANCJ is phosphorylated on serine 990. Formation of the BRCA1-FANCJ complex may be needed for the conversion of interstrand crosslinks (ICL) to DSB for repair by homologous recombination and to promote replication restart at G4 sites (35–38). Beyond these roles, the BRCT domain also binds to supercoiled DNA, contributes to chromatin remodeling, and participates in the detection of DNA breaks (39–42). It also suppresses ribosomal R-loops by facilitating sense-antisense rRNA pairing and enhancing RNA polymerase I-driven antisense rRNA transcription (43).

The central region of BRCA1, though unstructured, serves as a scaffold for protein and DNA interactions (23, 26, 42, 44). Within this region, two discrete DNA binding domains (DBD) have been identified: DBD1 (amino acids 330-554) and DBD2 (amino acids 894-1057) (40, 43, 45–47).

Although the relative affinities of the DBD for branched DNA substrates have been examined, the results were mostly qualitative due to the presence of multiple BRCA1-DNA complexes (23). To further characterize the DNA binding preferences of these domains, we have determined the affinities of DBD1, DBD2, and BRCT for three DNA substrates representative of repair intermediates: single-stranded DNA (ssDNA), double-stranded DNA (dsDNA), and G4 DNA. Our findings suggest that the BRCA1 DBD are functionally specialized, with distinct substrate specificities that may direct BRCA1 protein to either DSB or to the telomere during DNA damage response.

## MATERIALS AND METHODS

### Buffers and reagents

All solutions were prepared with analytical-grade chemicals and Type I ultrapure water from a Smart2Pure 6 UV/UF system (ThermoFisher; Waltham, MA, USA). The solutions were then sterilized through a 0.22 µm PES filter prior to use.

### Expression and purification of recombinant proteins

Codon-optimized *E. coli* expression plasmids encoding human BRCA1 DBD1 (aa330-554), DBD2 (aa894-1057), and BRCT (aa1646-1863) were purchased from VectorBuilder (Chicago, IL, USA). Constructs were transformed into NiCo21(DE3) competent cells (New England Biolabs; Ipswich, MA, USA). Starter cultures (125 mL) were expanded into 4 L of LB medium (1:20 dilution) that was maintained at 37 °C and 250 rpm using a MaxQ 4000 orbital shaker (ThermoFisher). Protein expression was induced with 1 mM IPTG when the OD_600_ reached 0.6. After 4 hours of expression, the cells were harvested by centrifugation using a J6-MI high-capacity centrifuge equipped with a JS-4.2 rotor (4,000 rpm at 4 °C for 90 minutes).

Cell pellets were resuspended in lysis buffer (20 mM NaPi (pH 7.5), 300 mM NaCl, 30 mM imidazole, 1 mM DTT, 5% (v/v) glycerol, 1% NP-40, and 1 mM PMSF), and lysed by sonication using a Fisherbrand 505 Dismembrator (Fisher Scientific; Hampton, NH, USA). Samples were chilled on ice during sonication (15 seconds on, 45 seconds off, and 45% amplitude for 15 minutes). Lysates were clarified by centrifugation with a Sorvall RC5C Plus high-speed centrifuge and a SS-34 fixed angle rotor (18,500 rpm at 4 °C for 90 minutes). The supernatant was filtered through a 0.45 µm PES membrane prior to liquid chromatography.

All protein purification steps were performed at 4 °C. Clarified lysates were incubated for 45 minutes with 20 mL Ni-NTA agarose resin (Goldbio; St. Louis, MO, USA) that was pre-equilibrated in binding buffer (20 mM NaPi (pH 7.5), 300 mM NaCl, 30 mM imidazole, 1 mM DTT, and 5% glycerol). After batch binding, the resin was washed sequentially with 10 column volumes (CV) of equilibration buffer and 10 CV of low-salt buffer (20 mM NaPi (pH 7.5), 30 mM NaCl, 30 mM imidazole, 1 mM DTT, and 5% glycerol). Bound proteins were eluted with 1 CV of elution buffer (20 mM NaPi (pH 7.5), 30 mM NaCl, 300 mM imidazole, 1 mM DTT, and 5% glycerol).

Eluates were immediately loaded onto a 5 mL HiTrap Heparin HP column (GE Healthcare, Chicago, IL, USA) that was equilibrated with 20 mM HEPES (pH 7.5), 30 mM NaCl, 5 mM TCEP, and 5% glycerol using an AKTA Start FPLC system (GE Healthcare). The column was washed with 10 CV of the equilibration buffer, and the protein was eluted with 20 mM HEPES (pH 7.5), 1 M NaCl, 5 mM TCEP, and 5% glycerol over a 50 mL linear gradient in 2 mL fractions. Elution fractions were analyzed by SDS-PAGE, pooled, and dialyzed into storage buffer (20 mM HEPES pH 7.5, 150 mM KCl, 5 mM TCEP, and 20% glycerol) using 3500 MWCO tubing (ThermoFisher). After three buffer exchanges, proteins were flash-frozen in liquid nitrogen and stored at −80 °C. Protein concentrations were measured with a NanoDrop One C spectrophotometer (ThermoFisher) using the molar extinction coefficients listed in Table 1.

**Table 1.**
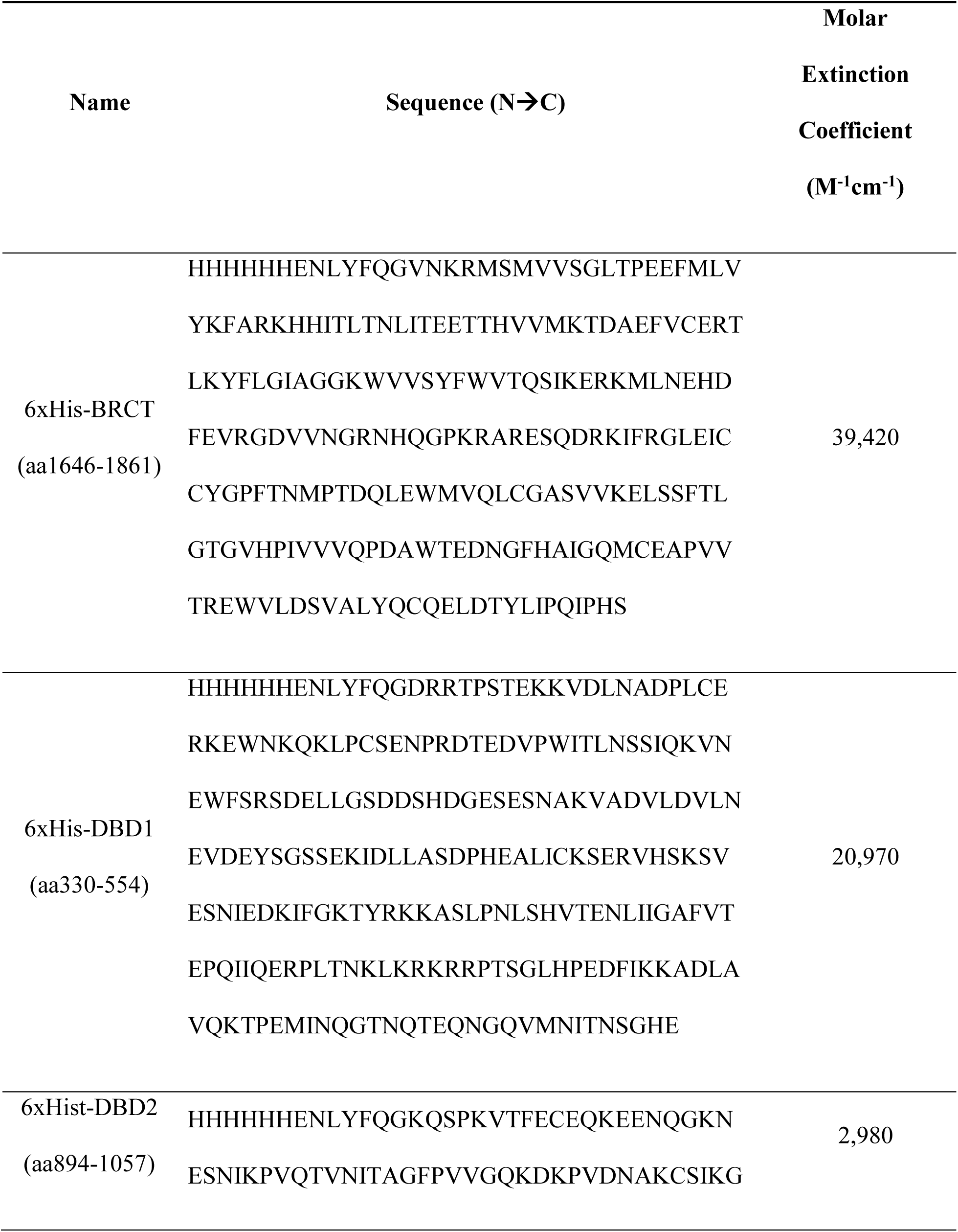
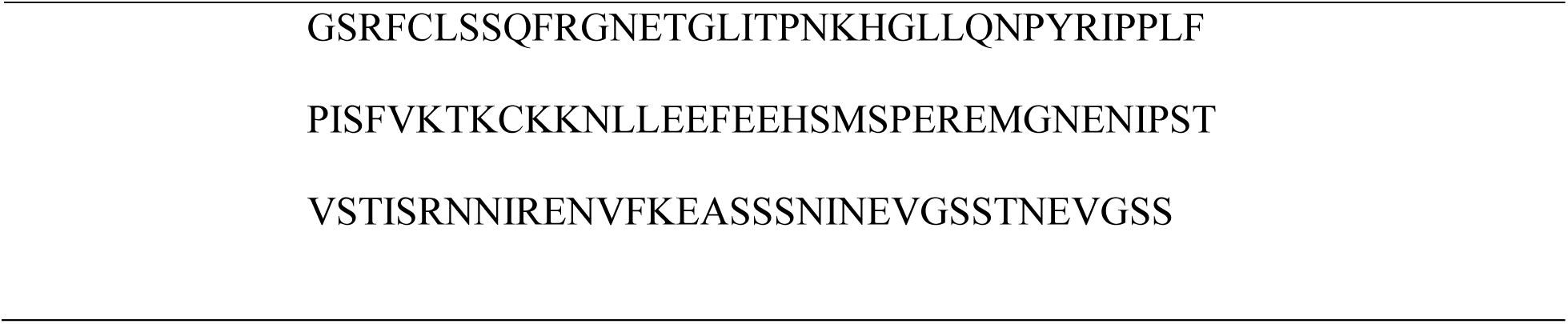
Amino acid sequences of BRCA1 DBD1, DBD2, and BRCT.

| Name | Sequence (N→C) | Molar<br>Extinction<br>Coefficient<br>(M<sup>-1</sup>cm<sup>-1</sup>) |
| --- | --- | --- |
| 6xHis-BRCT<br>(aa1646-1861) | HHHHHHENLYFQGVNKRMSMVVSGLTPEEFMLV<br>YKFARKHHITLTNLITEETTHVVMKTDAEFVCERT<br>LKYFLGIAGGKWVVS YFWVTQSIKERKMLNEHD<br>FEVRGDVVNGRNHQGPKRARES QDRKIFRGLEIC<br>CYGPFTNMPTDQLEWMVQLCGASVVKELSSFTL<br>GTGVHPIVVVQPD AWTEDNGFHAIGQMCEAPVV<br>TREWVLDSVALYQCQELDTYLIPQIPHS | 39,420 |
| 6xHis-DBD1<br>(aa330-554) | HHHHHHENLYFQGDRRTPSTEKKVDLNADPLCE<br>RKEW NKQKLPCSENPRDTE DVPWITLNSSIQKVN<br>EWFSRSD ELLGSDDSHDGESE SNAKVADVLDVLN<br>EVDEYSGSSEKIDLLASDPHEALICKSERVHSKSV<br>ESNIEDKIFGKTYRKKASLPNLSHVTENLIIGAFVT<br>EPQIIQERPLTNKLKRKR RPTSGLHPEDFIKKADLA<br>VQKTPEMINQGTNQTEQNGQVMNITNSGHE | 20,970 |
| 6xHis-DBD2<br>(aa894-1057) | HHHHHHENLYFQGKQSPKVTFECEQKEENQGKN<br>ESNIKPVQTVNITAGFPVVGQKDKPVDNAKCSIKG | 2,980 |

### DNA oligos

All oligonucleotides were synthesized by Integrated DNA Technologies (IDT; Coralville, IA, USA). Oligos were dissolved in Buffer H (20 mM HEPES pH 7.5, 150 mM KCl, 5 mM TCEP, and 5% glycerol), and DNA concentrations were determined spectrophotometrically. Double-stranded DNA was prepared by mixing the non-biotinylated complementary strand in 1.05x molar excess over the biotinylated oligo, incubating the sample at 95 °C for 10 minutes, and cooling it slowly to room temperature over 6 hours. A Cy3 label was included in the complementary strand to confirm successful annealing by gel electrophoresis.

### Biolayer interferometry (BLI)

BLI assays were performed on a BLItz instrument (Sartorius) in Buffer H at 25 °C. High precision streptavidin-coated biosensors (SAX) were purchased from Sartorius and hydrated in Buffer H for 10 minutes prior to use. For each experiment, the sensor was placed in Buffer H for 30 seconds to collect an initial baseline. Next, 150 nM of biotinylated DNA was loaded onto the sensor over 120 seconds and a second baseline was collected in Buffer H for 30 seconds. Protein samples were then introduced at different concentrations. The association reactions were monitored for 180 seconds, and dissociation of the complexes were examined for 120 seconds in Buffer H. A reference curve was obtained in buffer alone and was subtracted from the other traces. BLI measurements were performed in triplicates on separate days. For further methodological details, see reference (48).

### Data analysis

BLI binding isotherms were constructed by plotting the steady-state response (Δλ) values against protein concentration. The curves were fit to a 1:1 binding model in Scientist 3.0 software (Micromath; St. Louis, MO) using the following implicit formulas, where Δλ was the observed signal change, A was the amplitude, and K was the equilibrium association constant. *D_f_* and *D_t_* were the free and total concentration of DNA, while *P_f_* and *P_t_* described the free and total concentration of the protein. The binding constants reported were determined from the average and 95% confidence interval of three independent data sets.

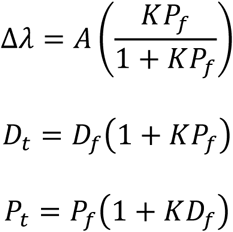

## RESULTS

### Expression and purification of BRCA1 DNA binding domains

Recombinant BRCA1 DBD1 (aa 330-554), DBD2 (aa 894-1057), and BRCT domain (aa 1646-1861) were expressed in *E. coli* and purified to near homogeneity using Ni-NTA and heparin affinity chromatography. All three proteins migrated as single bands at their expected molecular weights by SDS-PAGE (Figure 1).

**Figure 1.**
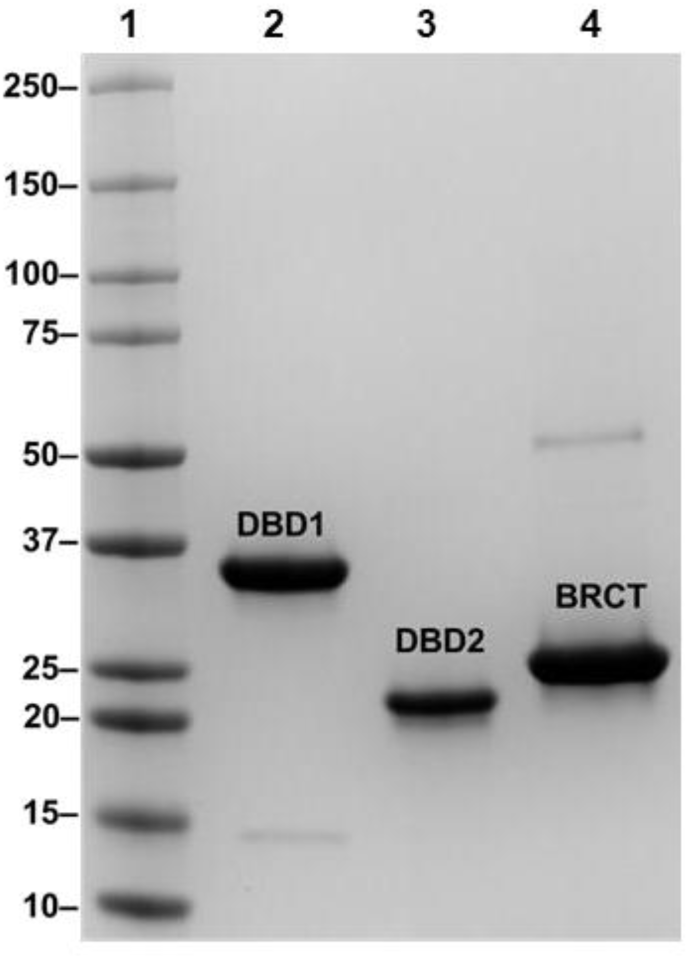
SDS-PAGE of purified recombinant BRCA1 DBD1, DBD2, and BRCT. A protein ladder with known molecular weights (kDa) was loaded into Lane 1. Purified DBD1 (Lane 2), DBD2 (Lane 3), and BRCT (Lane 4) were loaded in the respective lanes.

### Design of DNA substrates

To examine the DNA binding properties of the BRCA1 domains, we designed three biotinylated DNA substrates as shown in Table 2: a 60 nt ssDNA, a fully annealed blunt-end dsDNA, and a G4 formed from the human telomeric repeat sequence. The ssDNA and dsDNA structures are common repair intermediates in homologous recombination, while the human telomeric G4 was examined due to BRCA1’s role in telomere maintenance (28, 29). Branched DNA substrates were excluded maintain 1:1 binding stoichiometry for quantitative analysis of the affinity values.

**Table 2.**
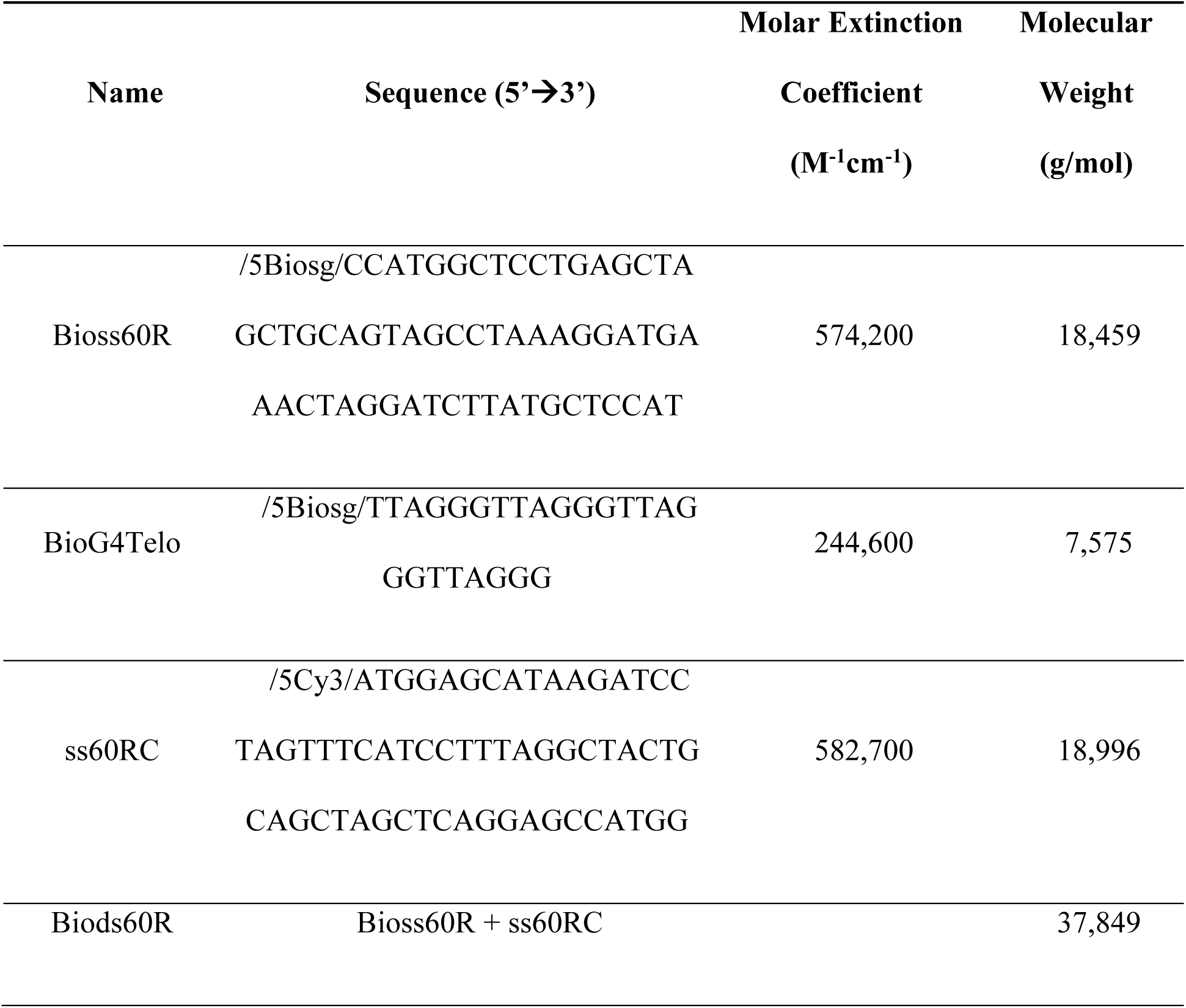
DNA oligo sequences.

### Biolayer interferometry assays

Biolayer interferometry (BLI) was used to measure the binding affinities of DBD1, DBD2, and BRCT for each DNA substrate. Figure 2A outlines a standard BLI workflow. The streptavidin-coated BLI sensors (SAX) were hydrated in Buffer G, and an initial baseline signal was measured (Phase 1). The biotinylated DNA was loaded onto the SAX biosensors, producing an increase in binding signal (Δλ) (Phase 2). After loading the DNA, the sensor tip was washed in Buffer G to establish a new baseline (Phase 3). This step ensured that the DNA substrate remain bound to the SAX sensors. The proteins were then added at varying concentrations to initiate the association reactions (Phase 4). After reaching a steady-state plateau, the sensor was placed into Buffer G to monitor the dissociation reactions (Phase 5). Figure 2B illustrates the association and dissociation time-courses collected from multiple protein concentrations. The plateau values were plotted against protein concentration and fit to a 1:1 interaction model using non-linear least squares analysis to determine the equilibrium constant (K) values (Figure 2C).

**Figure 2.**
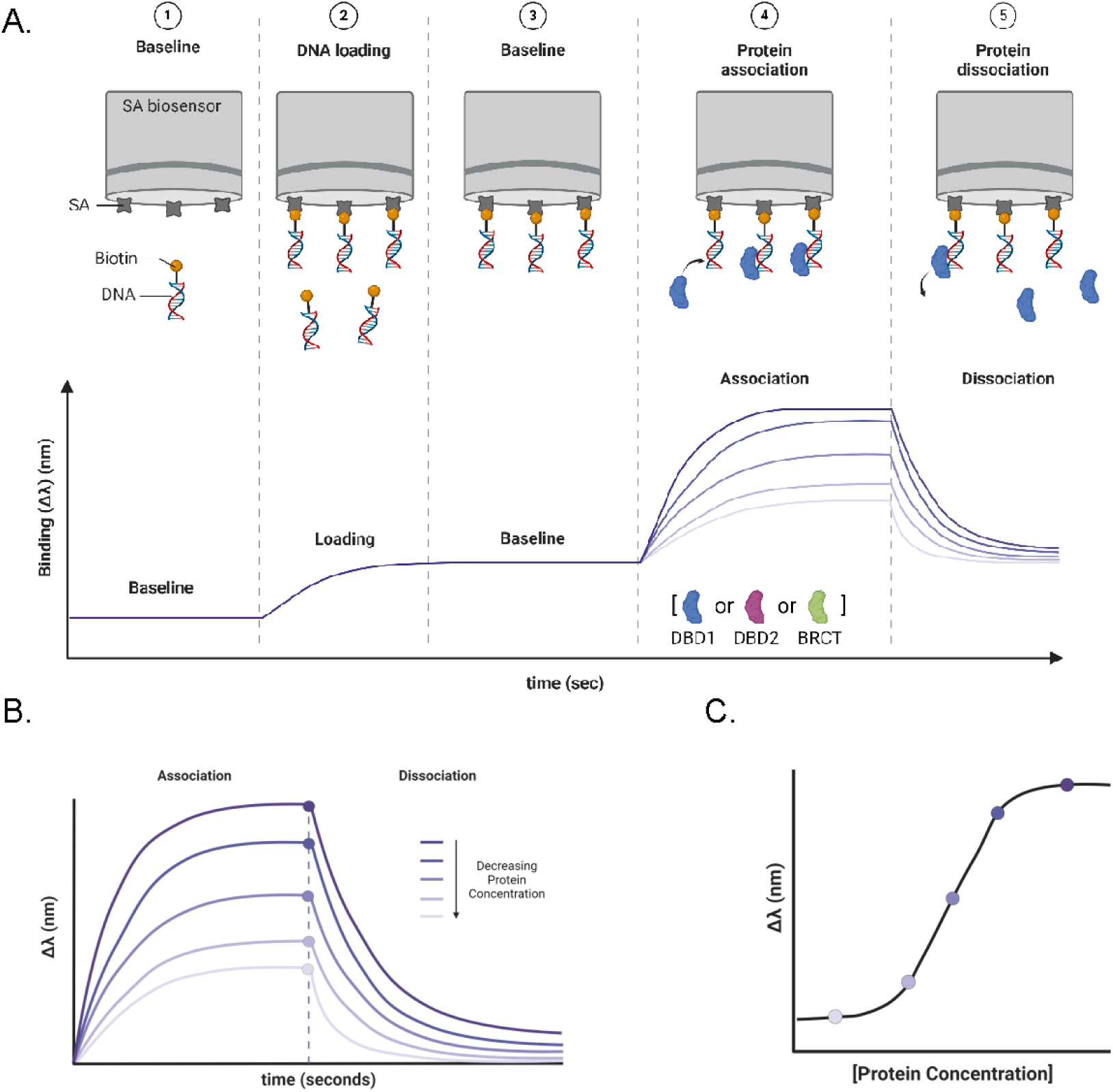
Biolayer interferometry (BLI) binding assays. **A.** Cartoon depicting a time-course of each step of a BLI experiment. The interference pattern of a streptavidin-coated (SAX) biosensor was measured in Buffer G to establish a baseline (Phase 1). Biotinylated DNA was loaded onto the sensor tip through biotin-SA interactions and an increase in Δλ was measured (Phase 2). The biosensor was washed with buffer to confirm the DNA substrate remained surface-immobilized and to establish a new baseline (Phase 3). Protein (DBD1, blue; DBD2, magenta; BRCT, green) was introduced to initiate the association reaction (Phase 4). Once a binding equilibrium was reached, the biosensor was returned to Buffer G to examine the dissociation of the protein-DNA complexes (Phase 5).

### DNA binding profiles of the BRCA1 domains

Quantitative BLI revealed distinct DNA substrate preferences for each BRCA1 domain. DBD1 exhibited the highest binding affinity for dsDNA (K = 2.92 ± 0.66 × 10^7^ M^-1^; K_D_ = 3.55 ± 0.90 × 10^-8^ M), followed by a lower affinity for ssDNA (K = 1.67 ± 0.65 × 10^7^; K_D_ = 6.84 ± 3.38 × 10^-8^ M), and the weakest affinity for G4 DNA (K = 1.52 ± 0.28 × 10^7^ M^-1^; K_D_ = 6.72 ± 1.25 × 10^-8^ M) (Figure 3). This preference for dsDNA is consistent with the potential role of DBD1 in recognizing DSB during early HR. Although DBD1 showed measurable binding to ssDNA and to G4 DNA, these interactions were weaker, indicating lower selectivity for these repair intermediates.

**Figure 3.**
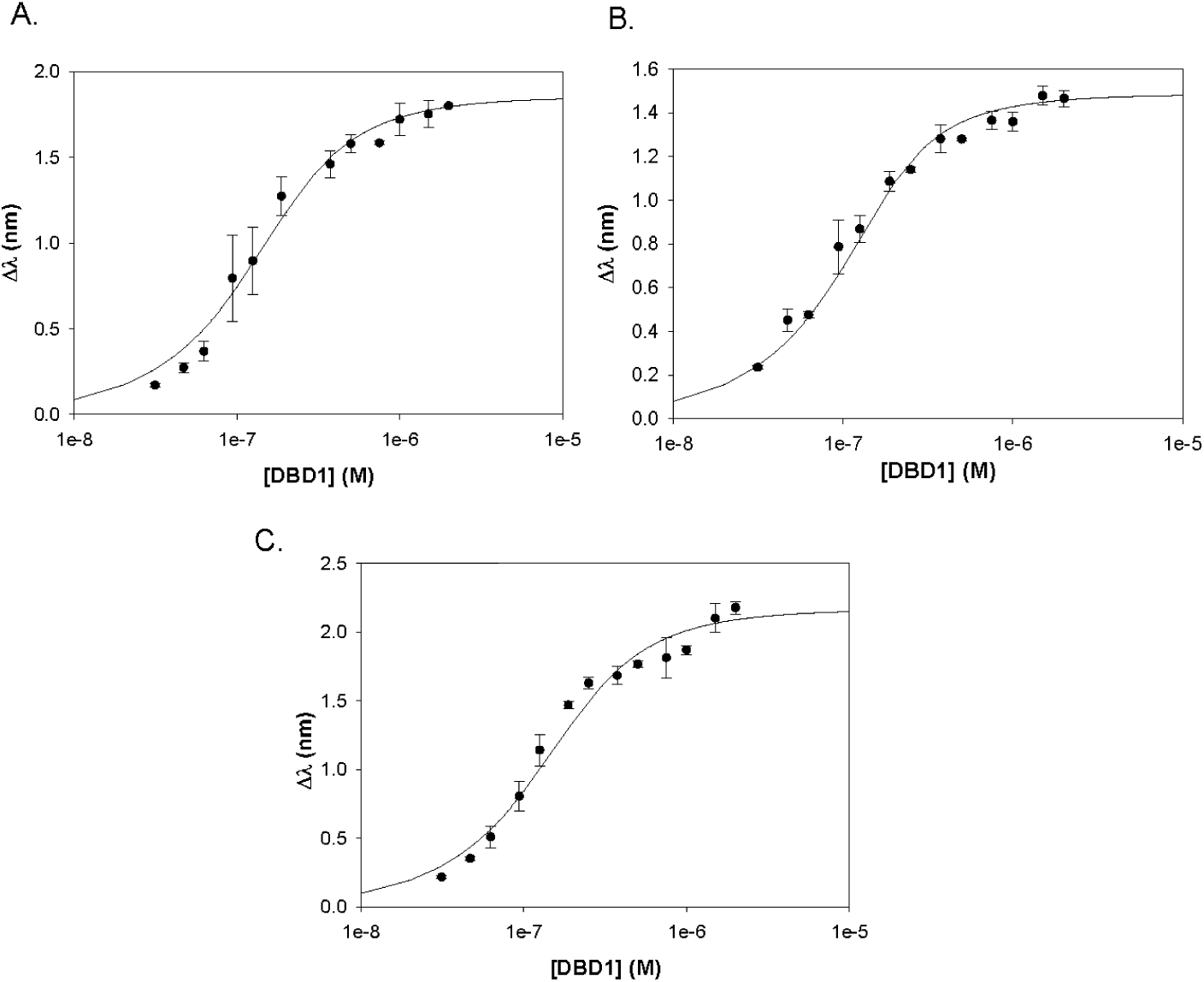
Plots of BRCA1 DBD1. A. ssDNA. B. dsDNA. C. G4 DNA.

DBD2 showed the opposite binding pattern, with the highest affinity for G4 DNA (K = 3.26 ± 1.21 × 10^7^ M^-1^; K_D_ = 3.47 ± 1.61 × 10^-8^ M), followed by ssDNA (K = 2.47 ± 0.23 × 10^7^ M^-1^; K_D_ = 4.08 ± 0.37 × 10^-8^ M), and the weakest binding to dsDNA (K = 2.27 ± 0.31 × 10^7^ M^-1^; K_D_ = 4.46 ± 0.60 × 10^-8^ M) (Figure 4). The enhanced preference for G4 structures suggests that DBD2 may contribute to BRCA1’s established role in resolving secondary structures at telomeres or fragile sites under replication stress.

**Figure 4.**
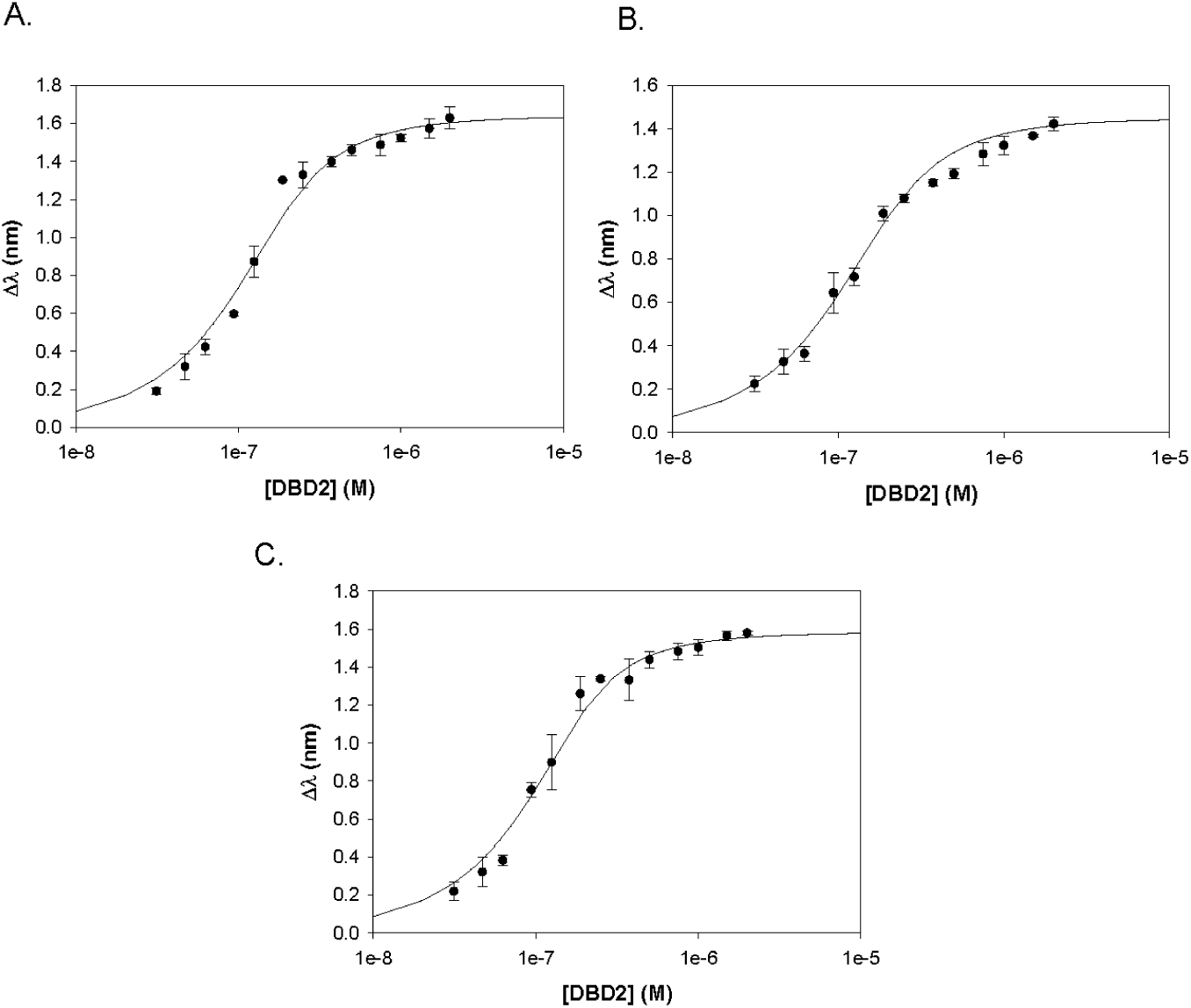
Plots of BRCA1 DBD2. A. ssDNA. B. dsDNA. C. G4 DNA.

The BRCT, although traditionally described as a phosphopeptide-binding domain, also demonstrated detectable binding to DNA substrates with the same affinity patterns as DBD2. The BRCT bound strongest to G4 DNA (K = 6.82 ± 1.09 × 10^6^ M^-1^; K_D_ = 1.49 ± 0.26 × 10^-7^ M), followed closely by ssDNA (K = 6.34 ± 0.49 × 10^6^ M^-1^; K_D_ = 1.58 ± 0.12 × 10^-7^ M), and the weakest binding to dsDNA (K = 5.46 ± 0.42 × 10^6^ M^-1^; K_D_ = 1.84 ± 0.14 × 10^-7^ M) (Figure 5). Notably, these affinity values were ∼10-fold lower than those observed for DBD2. All BLI binding results are summarized in Table 3.

**Figure 5.**
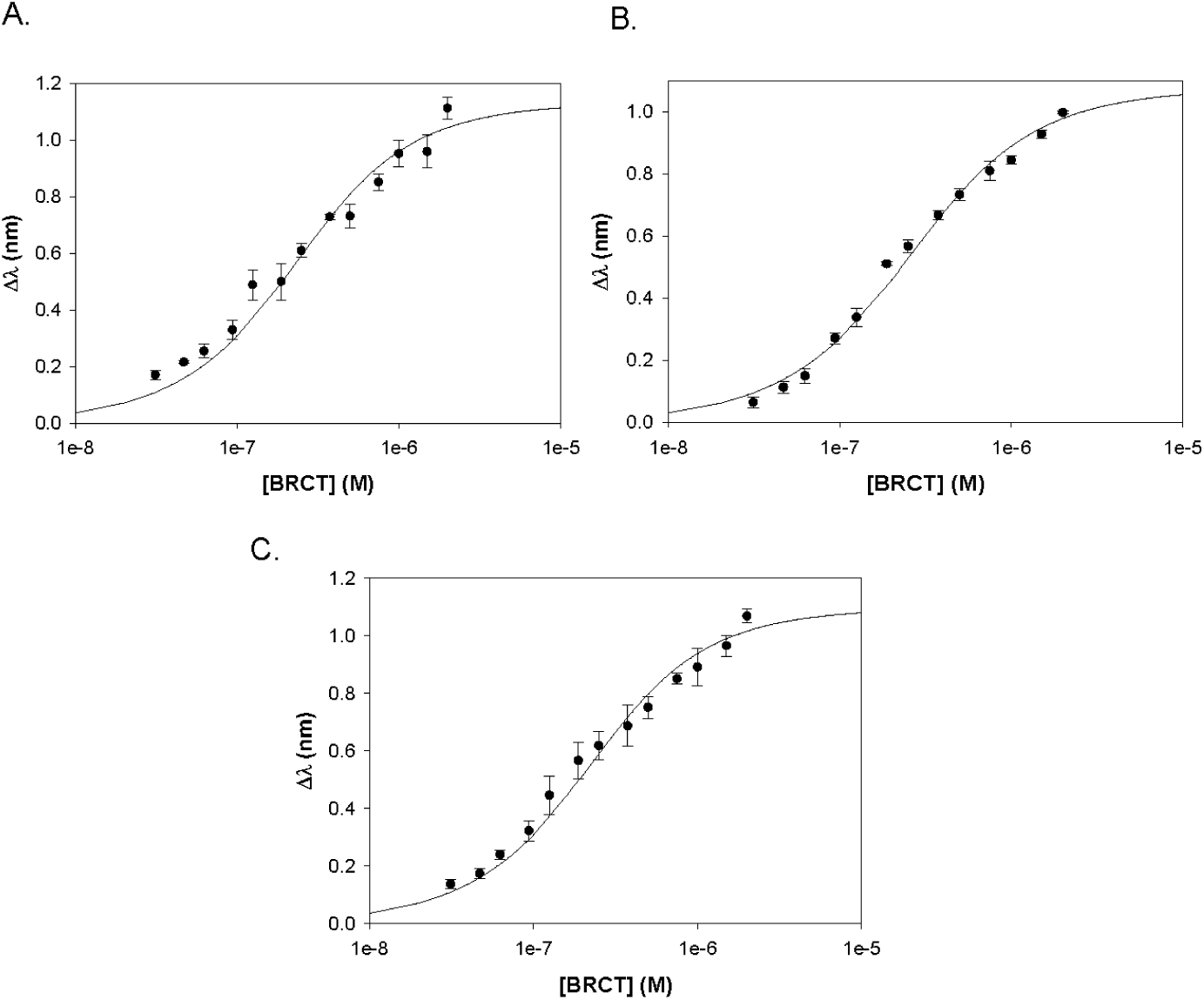
Plots of BRCA1 BRCT. A. ssDNA. B. dsDNA. C. G4 DNA.

**Table 3.** Summary of BLI data.

| Protein | DNA Substrate | K (M<sup>-1</sup>) | K<sub>D</sub> (M) | A | R<sup>2</sup> |
| --- | --- | --- | --- | --- | --- |
| **DBD1** | ssDNA | 1.67 ± 0.65 x 10<sup>7</sup> | 6.84 ± 3.38 x 10<sup>-8</sup> | 1.85 ± 0.037 | 0.995 |
|  | G4 | 1.52 ± 0.28 x 10<sup>7</sup> | 6.72 ± 1.25 x 10<sup>-8</sup> | 2.16 ± 0.044 | 0.995 |
|  | dsDNA | 2.92 ± 0.66 x 10<sup>7</sup> | 3.55 ± 0.90 x 10<sup>-8</sup> | 1.48 ± 0.021 | 0.997 |
| **DBD2** | ssDNA | 2.47 ± 0.23 x 10<sup>7</sup> | 4.08 ± 0.37 x 10<sup>-8</sup> | 1.64 ± 0.029 | 0.996 |
|  | G4 | 3.26 ± 1.21 x 10<sup>7</sup> | 3.47 ± 1.61 x 10<sup>-8</sup> | 1.58 ± 0.023 | 0.997 |
|  | dsDNA | 2.27 ± 0.31 x 10<sup>7</sup> | 4.46 ± 0.60 x 10<sup>-8</sup> | 1.45 ± 0.043 | 0.996 |
| **BRCT** | ssDNA | 6.34 ± 0.49 x 10<sup>6</sup> | 1.58 ± 0.12 x 10<sup>-7</sup> | 1.13 ± 0.027 | 0.993 |
|  | G4 | 6.82 ± 1.09 x 10<sup>6</sup> | 1.49 ± 0.26 x 10<sup>-7</sup> | 1.09 ± 0.021 | 0.995 |
|  | dsDNA | 5.46 ± 0.42 x 10<sup>6</sup> | 1.84 ± 0.14 x 10<sup>-7</sup> | 1.07 ± 0.016 | 0.997 |

## DISCUSSION

The BRCA1 central DNA binding domain (DBD) has been implicated in recognizing DNA structures that are critical to different repair pathways, including cruciform, supercoiled, G4, and triplex DNA (23, 26, 46, 47). However, definitions of the domain’s boundaries have varied. Paull et al. originally characterized the DBD as spanning residues 452–1079, while Brázda et al. refined this to aa 444–1057 (26), and Masuda et al. focused on aa 421–701 (25). Later studies suggested further subdivision—Mark et al. proposed two discrete subdomains (aa 498–663 and 936–1057), while Naseem et al. and Brázda et al. isolated minimal fragments (aa 340–554 and 894–1057, respectively) with DNA binding activity (44, 47, 49). Based on these studies, we generated BRCA1 constructs DBD1 (aa 330-554) and DBD2 (aa 894-1057) to clarify the specific DNA substrate preferences of these domains.

In addition, we examined the BRCT domain (aa 1646-1861), a structured phosphoprotein-binding domain with emerging roles in DNA and chromatin interactions (50). While BRCT has been studied extensively in the context of protein-protein interactions, its direct contributions to DNA binding remained less defined. Importantly, prior studies did not provide DNA binding affinity values of the isolated BRCA1 domains, nor did they assess whether the individual domains acted independently or synergistically to engage specific DNA targets.

Using BLI, we found that DBD1 preferentially binds to dsDNA with highest affinity, followed by ssDNA and G4 DNA. In contrast, DBD2 and BRCT both exhibited strongest binding to G4 DNA, intermediate binding to ssDNA, and the weakest binding to dsDNA. These findings reflect a functional division of labor among the BRCA1 domains and suggest specialization to form distinct repair intermediates. The 1:1 stoichiometry observed in our BLI experiments supports a model in which these domains engage with DNA substrates directly and independently. DBD1’s preference for dsDNA implies a primary role in early HR, potentially acting during the recognition of the blunt DSB ends and the recruitment of nucleases for end resection (7–11). DBD2’s preference for G4 DNA is consistent with previous reports linking BRCA1 to the resolution of replication stress and telomeric structures, where G4 DNA commonly form (28, 29). Interestingly, BRCT’s lower but selective affinity for G4 and ssDNA supports its proposed role in chromatin remodeling and transcriptional regulation at G4-prone loci (41,43).

These binding preferences are further supported by structural models. As shown in Figure 6, DBD1 and DBD2 are predicted to occupy opposite, solvent-accessible surfaces of BRCA1, consistent with functional specialization. DBD1 is more surface-exposed and is likely accessible to dsDNA at DSB sites. DBD2 and BRCT, on the other hand, are colocalized on the same surface and may coordinate binding to non-canonical B-form DNA such as G4 structures at the telomeres and chromatin. Since the BRCT is positioned internally within the full-length BRCA1 model, it will have less access to the DNA and thus has lower binding affinities compared to the DBD. DBD2 and BRCT may function together in binding to their DNA targets due to their similar positioning on the left-most side of the model. A potential explanation for the difference in binding preferences is that the domains undergo structural changes when bound to DNA. DBD1 and DBD2 are primarily unstructured domains, while BRCT is composed of α helix and β sheets (51). However, circular dichroism studies (see Supporting Information) observed no significant changes in secondary structure contact when any of the domains were bound to DNA. It remains possible that the full-length BRCA1 protein undergoes large-scale conformational rearrangements to accommodate or stabilize binding to specific DNA structures, particularly within multiprotein complexes.

**Figure 6.**
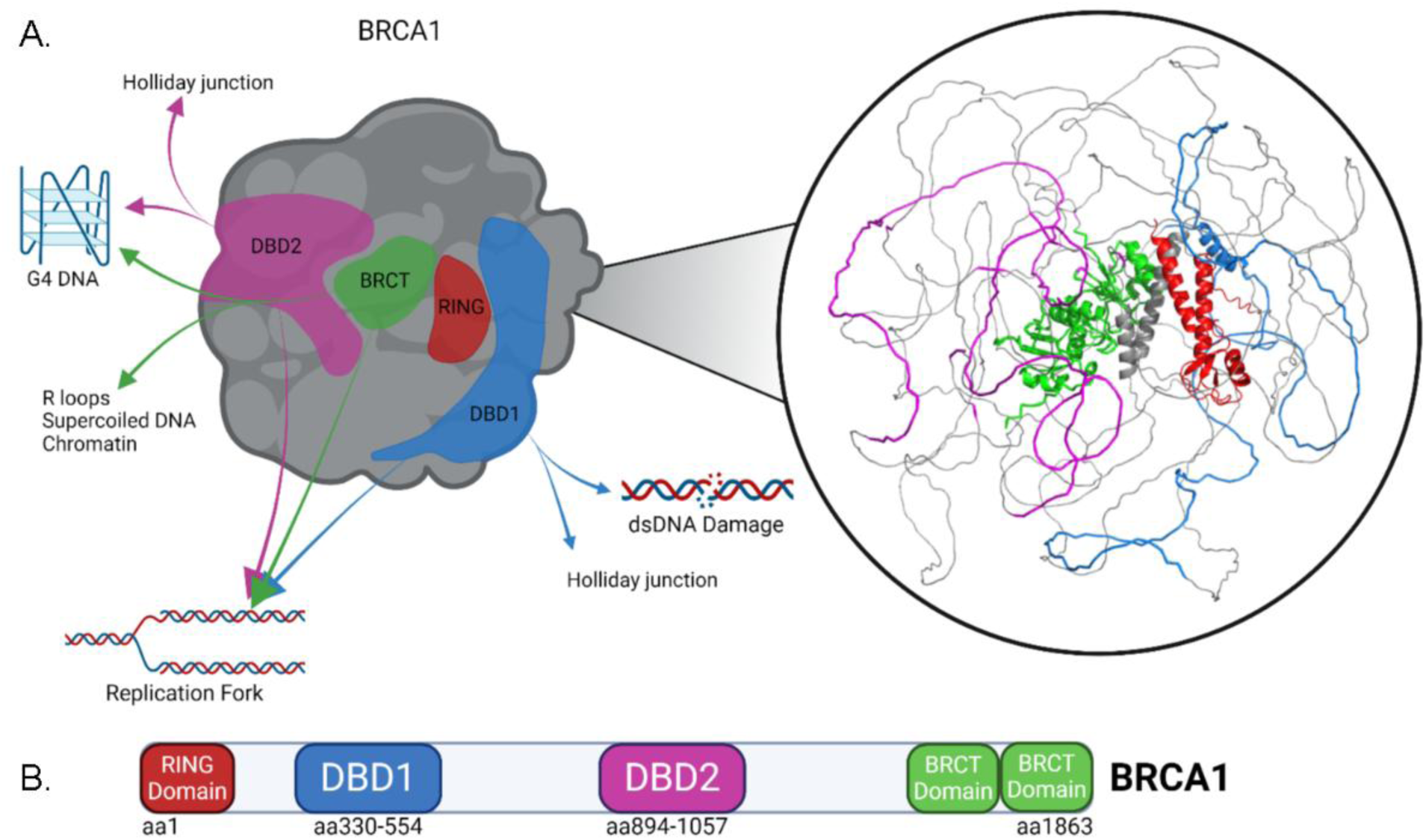
Models and map of BRCA1. **A.** Structural prediction of full-length BRCA1 using AlphaFold2 (right). Cartoon model (left) depicts the preferences of the different regions within BRCA1 in binding to distinct DNA structures. **B.** Cartoon map of BRCA1, highlighting domains RING (aa1-109, red), DBD1 (aa330-554, blue), DBD2 (aa894-1057, magenta), and the tandem BRCT repeats (aa1646-1863, green).

Our results both confirm and refine prior findings. For example, Masuda et al. observed that residues 421–701 prefer dsDNA over ssDNA (25), aligning with the behavior of our DBD1 construct (aa 330–554), which overlaps this region. In contrast, Sturdy et al. reported that aa 230–534 preferentially bind four-way junctions (46), which may reflect DBD1 recognizing each arm of the cruciform as individual dsDNA sites. This highlights the importance of substrate design when interpreting DNA binding profiles. Similarly, the broader DBD2-containing fragment (aa 444–1057) used by Brázda et al. exhibited strong binding to G4 and cruciform DNA, a trend preserved in our DBD2 construct (26). These comparisons suggest that DBD2 is the primary determinant of BRCA1’s affinity for non-canonical DNA structures, even within larger central constructs. We also observed BRCT binding to dsDNA, consistent with studies by Yamane et al. and Matsumoto et al. (43, 52), and further support the model proposed by Hu et al. in which BRCT contributes to chromatin remodeling at telomeres (41). Given that our G4 was derived from human telomeric repeat sequences, the BRCT’s interactions with this substrate further suggests BRCA1’s involvement in telomeric surveillance.

## CONCLUSIONS

Our findings provide quantitative evidence that the BRCA1 DBD exhibit distinct substrate preferences, with DBD1 favoring dsDNA and DBD2 and BRCT preferentially targeting G4 and ssDNA. These data support a functional model in which BRCA1 is guided to specific genomic sites based on domain-specific recognition of DNA structure—DBD1 facilitates BRCA1’s recruitment to DSB during early stages of HR; DBD2 and BRCT together enable BRCA1 localization to G4-rich regions such as telomeres or stalled replication forks, where BRCA1 helps coordinate chromatin remodeling and replication restart. Further studies are needed to determine whether the DBD1-2 fusion (aa 330-1057) exhibits additive or cooperative binding behavior and whether pathogenic BRCA1 variants within these domains alter DNA binding affinity or substrate specificity. A detailed understanding of these BRCA1-DNA interactions may clarify how specific mutations contribute to cancer risk and guide interpretation of variants of unknown significance (VUS) in clinical studies.

## ASSOCIATED CONTENT

## Supporting Information

The following files are available free of charge.

Circular dichroism (CD) spectroscopy data (PDF)

## ACCESSION CODES

BRCA1, UniProt: P38398

## AUTHOR INFORMATION

## Author Contributions

C.G.W. designed experiments. K.L., L.C., and C.G.W. acquired data and interpreted results. K.L., L.C., E.C., and C.G.W. prepared the manuscript draft. All authors have given approval to the final version of the manuscript.

## Funding Sources

This work was supported by AHA grant 19AIREA34460026 to C.G.W.

## ACKNOWLEDGMENT

Figures 2 and 6, as well as the abstract graphic, were created with BioRender.com. We thank the Spies Lab for providing the BRCA1 BRCT construct.

## ABBREVIATIONS

*BRCA1*: breast cancer susceptibility gene 1
HR: homologous recombination
DSB: double-strand break
DBD: DNA binding domain
aa: amino acid
BRCT: BRCA1 C-terminal repeats
ssDNA or Bioss60R: single-stranded DNA
dsDNA or Biods60R-Cy3: blunt-end double-stranded DNA
G4 DNA or BioG4Telo: G-quadruplex DNA
53BP1: tumor suppressor p53-binding protein 1
PALB2: partner and localizer of BRCA2
BRCA2: breast cancer type 2 susceptibility protein
RAD51: DNA repair protein RAD51 homolog 1
NHEJ: non-homologous end-joining
RING: really interesting new gene finger domain
FANCJ: Fanconi anemia complementation group J
BLI: biolayer interferometry
SA: streptavidin

---

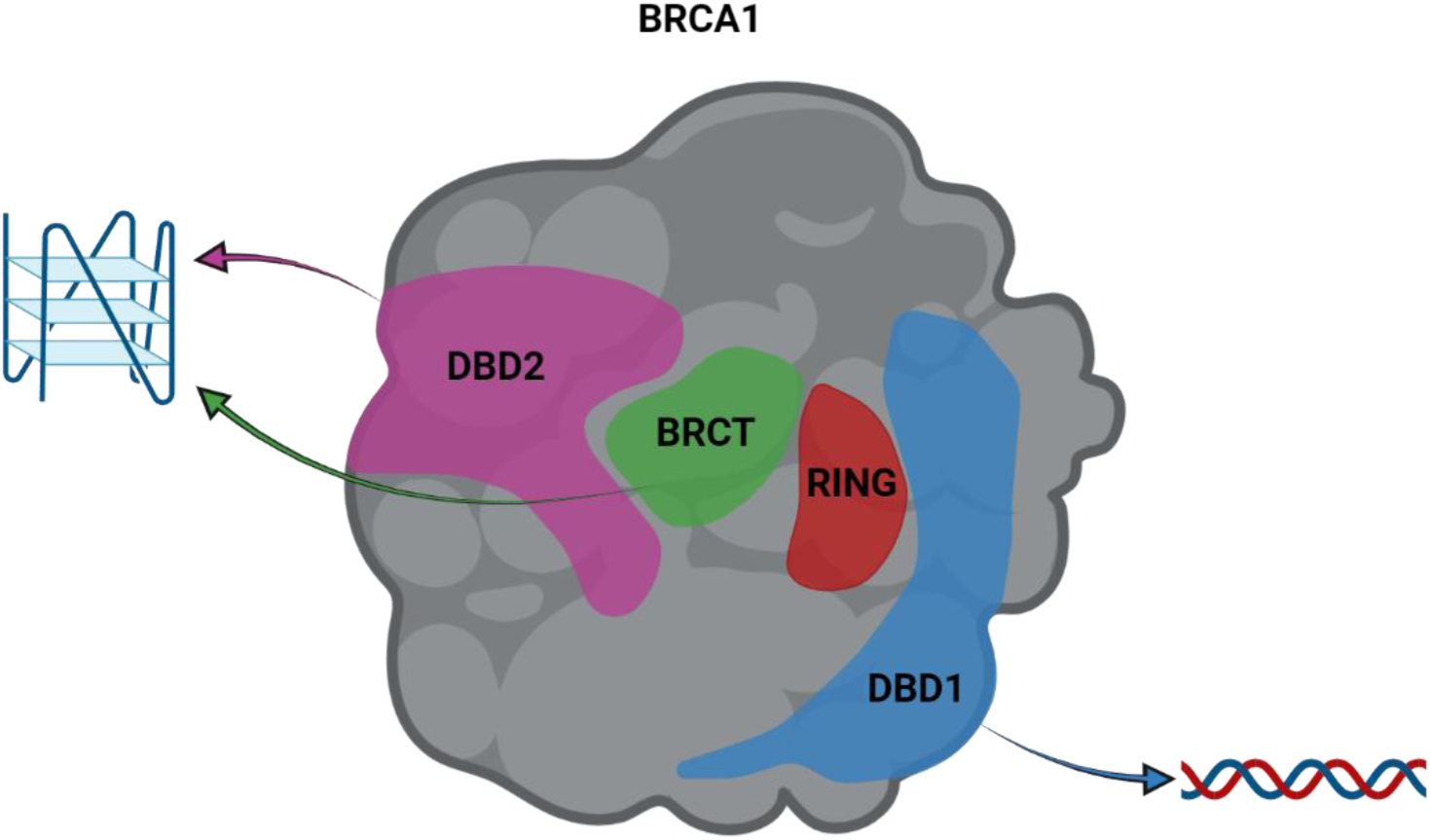

For Table of Contents only.

## Notes

### Competing Interest Statement

The authors have declared no competing interest.

### Summary of Updates

Fixed additional minor typos and grammatical errors.

