## Supporting Figure 1 for "Domain-specific DNA binding activities of BRCA1 reveal substrate preferences for homologous recombination and telomere regulation"

**Circular dichroism (CD) spectroscopy.** CD data was acquired using a JASCO J-815 spectropolarimeter (JASCO Inc.; Easton, MD, USA) equipped with a PTC-423S Peltier temperature control system. Protein and DNA samples were dialyzed into a buffer containing 20 mM Tris HCl (pH 7.5), 150 mM KCl, 1 mM DTT, and 5% glycerol. CD measurements were recorded at 25 °C across a wavelength range 200-260 nm for the BRCA1 DBD1 and DBD2. DNA substrates were titrated into the protein samples to assess changes in secondary structure upon binding. For each sample, five scans were collected and averaged. Reference spectra of buffer and DNA alone were subtracted from the averaged signal to obtain the final data.

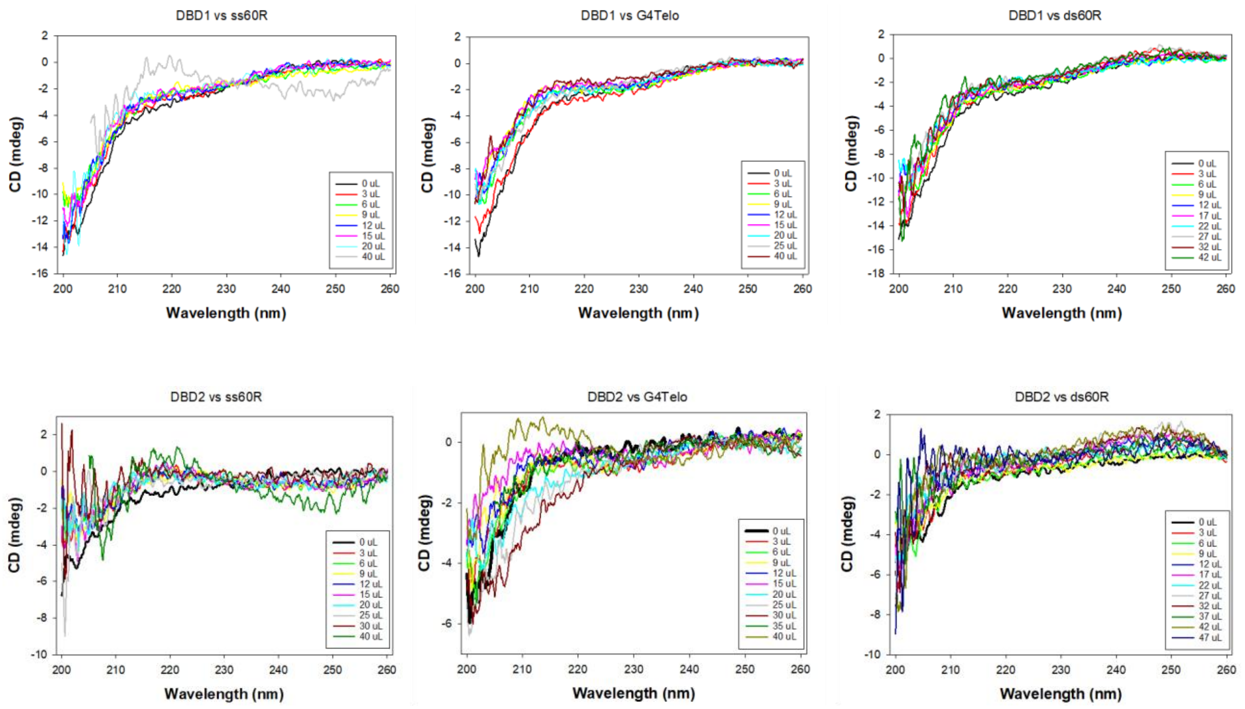


**Supporting Figure 1**. Circular dichroism spectra of DBD1 and DBD2 as a function of ssDNA, dsDNA, and G4 added. The results indicate that the domains remain intrinsically disordered upon binding to the DNA substrates.
